# Ketone Catabolism is Essential for Maintaining Normal Heart Function During Aging

**DOI:** 10.1101/2025.03.18.643760

**Authors:** Mariko Aoyagi Keller, Michinari Nakamura

**Affiliations:** Department of Cell Biology and Molecular Medicine, Rutgers New Jersey Medical School, Newark, New Jersey, 07103, USA

**Keywords:** ketone, ketolysis, SCOT, ketone catabolism, heart failure, ketogenic diet, hypertrophy, ketone oxidation, cardiac remodeling

## Abstract

The heart utilizes various nutrient sources for energy production, primarily favoring fatty acid oxidation. While ketones can be fuel substrates, ketolysis has been shown to be dispensable for heart development and function in mice. However, the long-term consequences of ketolysis downregulation in the heart remain unknown. Here we demonstrate that ketone catabolism is essential for preserving cardiac function during aging. The cardiac expression of succinyl-CoA:3-ketoacid CoA transferase (SCOT), a rate-limiting enzyme in ketolysis, decreases with aging in female mice. SCOT cardiomyocyte-specific knockout (cKO) mice exhibit normal heart function at 10 weeks of age but progressively develop cardiac dysfunction and remodeling as they age, without overt hypertrophy in both sexes. Notably, ketone supplementation via a ketogenic diet partially rescues contractile dysfunction in SCOT cKO mice, suggesting ketone oxidation-independent mechanisms contribute to the development of cardiomyopathy caused by SCOT downregulation. These findings indicate that ketone catabolism is crucial for maintaining heart function during aging, and that ketones confer cardioprotection independently of ketone oxidation.

## INTRODUCTION

The heart utilizes various nutrients, including fatty acids, glucose, ketones, amino acids, and lactate, to sustain a continuous and substantial energy supply (1). Although long-chain fatty acids serve as the primary energy substrates (2), the ability of the heart to flexibly utilize multiple nutrients acts as a compensatory mechanism that enables uninterruptedly pumping throughout life (3, 4). Indeed, mice with a genetic deletion of PPARα or CD36, key regulators of fatty acid metabolism, are viable and exhibit normal cardiac function, accompanied by a compensatory increase in the rates of glucose oxidation and glycolysis (5–7). Conversely, the genetic deletion of GLUT4, a key transporter in glucose metabolism, leads to severe cardiac hypertrophy and premature death in mice (8). Meanwhile, mice with cardiac-specific knockout (cKO) of GLUT4 develop modest cardiac hypertrophy but maintain normal contractile function without cardiac fibrosis, exhibiting normal serum concentrations of insulin, glucose, and fatty acids, as well as a normal lifespan (9). These findings highlight the metabolic flexibility of the heart, enabling it to utilize a variety of available nutrients for energy production. However, PPARα KO mice exhibit an accelerated cardiac aging phenotype, characterized by myocardial fibrosis and abnormal mitochondrial ultrastructure (10), whereas inhibition of CD36 attenuates age-associated cardiac decline by reducing intramyocardial lipid accumulation (11), underscoring a compensatory mechanism that employs diverse nutrients to limit cardiac aging.

Succinyl-CoA:3-ketoacid CoA transferase (SCOT) plays a pivotal role in ketone metabolism by catalyzing the transfer of coenzyme A from succinyl-CoA to acetoacetate. Mutations in the *3-oxoacid CoA-transferase 1* (*OXCT1*) gene, which encodes SCOT, lead to SCOT deficiency syndrome, characterized by recurrent ketoacidosis attacks (12). Systemic deletion of OXCT1 in mice exhibits normal prenatal development but causes neonatal lethality due to hyperketonemic hypoglycemia (13). Mice with a cardiac-specific genetic deletion of SCOT are viable, with normal body size, systemic metabolism, and fertility (14), but exhibit exacerbated oxidative stress, hypertrophy, and heart failure in response to transverse aortic constriction (15), suggesting that ketone oxidation is crucial for maintaining redox homeostasis and normal structure and function during pressure overload. Given that failing hearts upregulate ketone metabolism (16, 17) and can utilize ketones as fuel to improve cardiac efficiency, though with inconsistent findings (18–20), increased ketone utilization is generally considered an adaptive response (21). Consistent with these observations, we demonstrated that ketone supplementation through a ketogenic diet attenuates cardiac hypertrophy and remodeling in response to pressure overload (22). Furthermore, a ketogenic diet increases lysine acetylation in the heart (22), with acetylation of lysine acetyltransferase 6A (KAT6A) at K815 identified as a crucial regulator of cardiomyocyte size and mitochondrial function through its physical and functional interaction with AMPK (23, 24). These findings emphasize the role of ketones as signaling molecules (25), in part through modulating acetylation reactions (24), rather than merely oxidative substrates. However, it remains unknown whether ketone metabolism is essential for maintaining mechanical and structural cardiac integrity during aging.

In this paper, we investigate whether ketolysis influences cardiac function and morphology in aged mice. Here, we demonstrate that ketone catabolism is essential for maintaining normal cardiac function during aging. Furthermore, ketone supplementation via a ketogenic diet attenuates the development of dilated cardiomyopathy phenotypes induced by SCOT cardiac deletion in both sexes in mice, suggesting that ketones exert a ketone oxidation-independent cardioprotective mechanism.

## RESULTS

### Cardiac-specific SCOT knockout mice progressively develop contractile dysfunction in males

First, we examined whether SCOT expression in the heart changes with aging. SCOT expression levels were assessed in female mouse hearts at 20 days (pre-weaning, neonatal), 6 weeks (adolescent), and 8 months (middle-aged). In adolescent/young mice, SCOT was highly expressed in the heart, whereas its expression was suppressed before weaning and progressively declined with aging (Fig. 1a, 1b). To investigate the functional role of SCOT in the heart, we generated SCOT cKO mice by crossing *OXCT1* floxed mice (obtained from KOMP, Oxct1^tm1c^(KOMP)Wtsi) with αMHC-Cre mice (Fig. 1c). Successful knockdown of SCOT in the heart was confirmed by immunoblot analysis (Fig. 1d). BDH1, another ketolysis enzyme responsible for converting from β-hydroxybutyrate (β-OHB) to acetoacetate in the heart, was slightly upregulated in SCOT cKO mouse hearts, presumably as a compensatory response to ketolysis downregulation by SCOT deletion.

**Figure 1:**
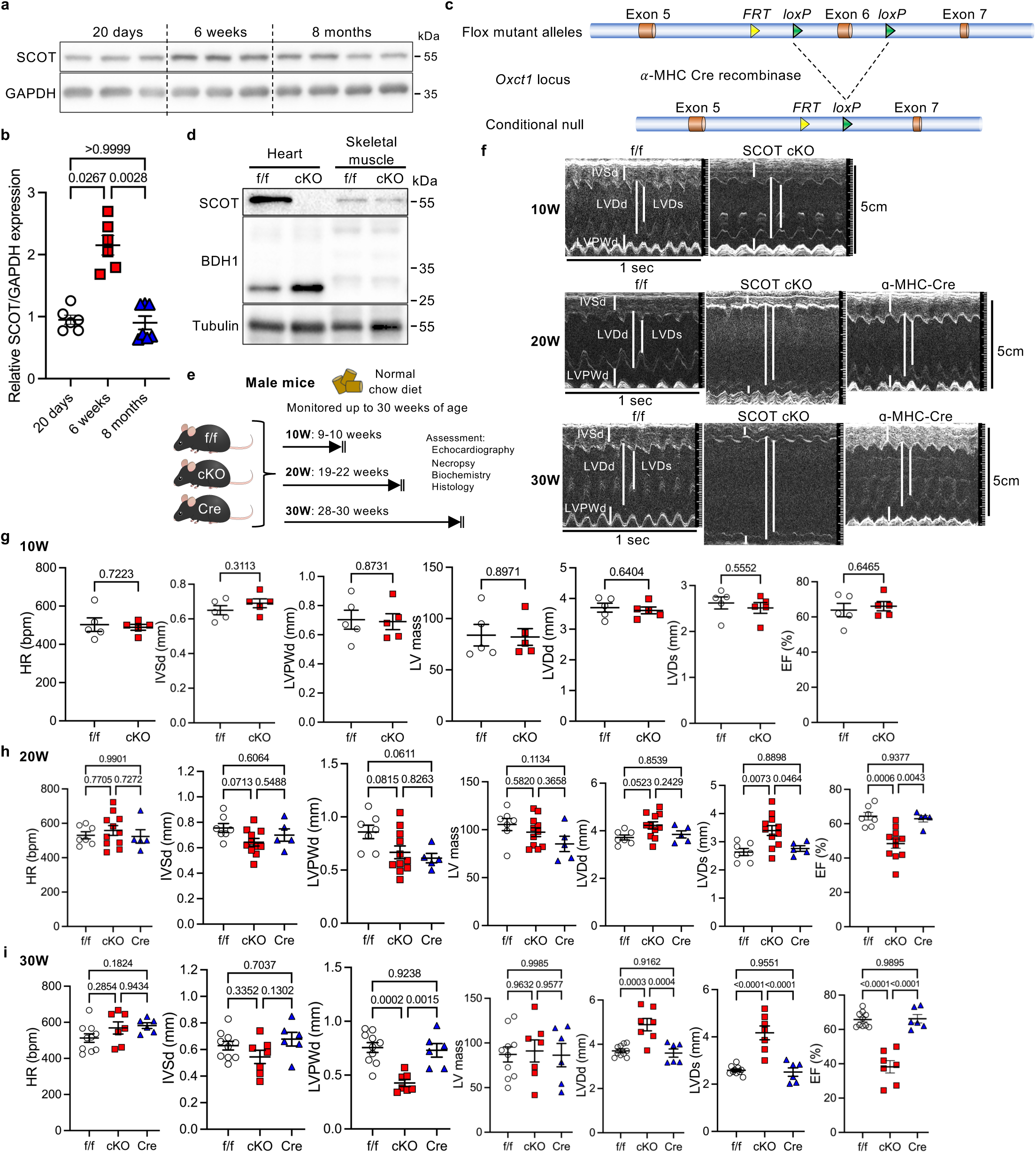
Cardiac-specific deletion of SCOT induces cardiac contractile dysfunction without overt hypertrophy in male mice. **a**, Immunoblots showing SCOT expression in the hearts of female mice at the indicated ages. **b**, Quantification of SCOT expression. n = 6 (20 days and 6 weeks), n = 7 (8 months). **c**, Schematic representation of the *OXCT1* gene-targeting strategy. **d**, Immunoblots confirming cardiac-specific knockout (cKO) of SCOT. **e**, Schematic illustration of the experimental design. Both floxed and αMHC-Cre alone mice were used as controls in the 20W and 30W age groups. **f**, Representative M-mode echocardiography images. Vertical scale bar, 5mm; transverse scale bar, 1 sec. **g-i**, Echocardiographic parameters of male mice at the indicated ages. n = 5 (g); n = 7 (f/f), 11 (cKO), and 5 (Cre) (h); n = 10 (f/f), 7 (cKO), and 6 (Cre) (i). One-way ANOVA followed by Tukey’s multiple comparison test. *N* represents biologically independent replicates. *p* values are indicated in the figure. Data are presented as mean ± SEM. Source data are provided as a Source Data file.

As reported previously (14), SCOT cKO mice were viable. Since prior studies did not examine the age-associated effects of SCOT downregulation on cardiac function (14, 15), we monitored SCOT cKO male mice for up to 30 weeks (Fig. 1e). To exclude potential effects of the αMHC-Cre transgene (26), two control groups (homozygous floxed (f/f) without αMHC-Cre and αMHC-Cre alone) were included in the long-term follow-up experiments. In agreement with previous findings, wall thickness and contractile function, as assessed by echocardiography, were comparable between SCOT cKO and f/f control male mice at 9-10 weeks of age (Fig. 1f, 1g). However, systolic function gradually declined by 19-22 weeks of age (Fig. 1f, 1h), and by 28-30 weeks, cardiac dilatation and reduced posterior wall thickness were observed (Fig. 1f, 1i).

Consistent with these observations, left ventricular (LV) mass, as assessed by echocardiography, remained unchanged between groups during aging (Fig. 1i). However, contractile function continued to deteriorate at later time points (Fig. 1i), indicating that SCOT downregulation in the heart progressively induces systolic dysfunction and cardiac remodeling with minimal impact on cardiac hypertrophy. These findings highlight the crucial role of SCOT expression in maintaining normal cardiac function.

### Heart weight increases following cardiac remodeling in SCOT cKO mice, indicating a secondary effect of heart failure

Since impaired systolic function was observed in SCOT cKO mice after 19-22 weeks of age, we further evaluated these mice through necropsy analysis (Fig. 2). Body weight and spleen weight remained comparable among the three groups throughout aging (Fig. 2a, 2b). Although ventricular weight normalized to tibia length was similar at 19-22 weeks of age (Fig. 2a, 2b), it was slightly increased in SCOT cKO mice at 28-30 weeks (Fig. 2d), supporting the notion that cardiac hypertrophy is not the primary cause of contractile dysfunction. Lung weight was significantly increased in cKO mice at 28-30 weeks (Fig. 2d), suggesting lung congestion due to heart failure. In contrast, liver weight was reduced in αMHC-Cre mice at the same time point (Fig. 2d). Gross appearance of the heart showed enlargement of the heart in SCOT cKO mice (Fig. 2b, 2e), indicating progressing cardiac remodeling following SCOT deletion. Since SCOT catalyzes the transfer of coenzyme A from succinyl-CoA to acetoacetate, we examined whether long-term deletion of cardiac SCOT affects serum β-OHB levels. However, cardiac SCOT deletion did not alter serum β-OHB levels throughout aging (Fig. 2c, 2f). These findings suggest that SCOT deletion induces cardiac remodeling and contractile dysfunction over time, independently of cardiac hypertrophy, and without affecting systemic ketone metabolism.

**Figure 2:**
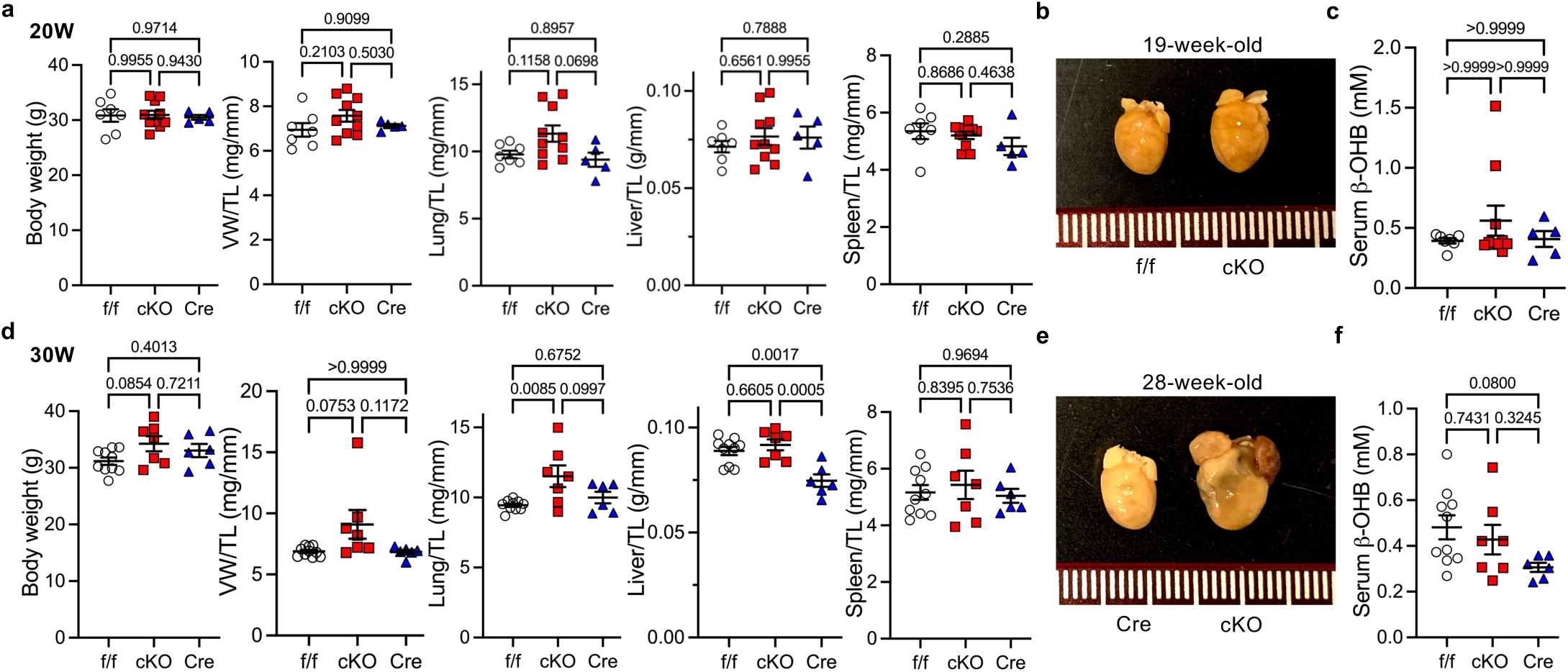
Cardiac-specific deletion of SCOT induces cardiac dilatation without overt systemic effects in male mice. a-c, Male mice at 19-22 weeks of age. n = 7 (f/f), 10 (cKO), and 5 (Cre). **d-f**, Male mice at 28-30 weeks of age. n = 10 (f/f), 7 (cKO), and 6 (Cre). Necropsy data, including body weight, ventricular weight (VW) normalized to tibia length (TL), lung weight/TL, liver weight/TL and spleen weight/TL (a, d). Gross heart morphology of the indicated mice (b, e). Scale unit: mm. Serum β-hydroxybutyrate levels (c, f). One-way ANOVA followed by Tukey’s multiple comparison test. *N* represents biologically independent replicates. *p* values are indicated in the figure. Data are presented as mean ± SEM. Source data are provided as a Source Data file.

Heart weight increased after systolic dysfunction developed in SCOT cKO mice, suggesting that this increase may be secondary to cardiac remodeling and heart failure. To confirm this, we assessed individual cardiomyocyte size using wheat germ agglutinin (WGA) staining. This analysis revealed that individual cardiomyocyte size remained comparable between f/f and SCOT cKO mice throughout aging (Fig. 3a, 3b), indicating no or minimal direct effect of SCOT deletion on cardiomyocyte size. Interestingly, αMHC-Cre mice exhibited smaller cardiomyocyte size at 28-30 weeks of age (Fig. 3b). Furthermore, we assessed cardiac fibrosis using Picric Acid Sirius Red staining and found significantly increased fibrosis in SCOT cKO mouse hearts as early as 19-22 weeks of age (Fig. 3c, 3d). These results suggest that SCOT plays a crucial role in maintaining contractile function and regulating inflammation in the heart.

**Figure 3:**
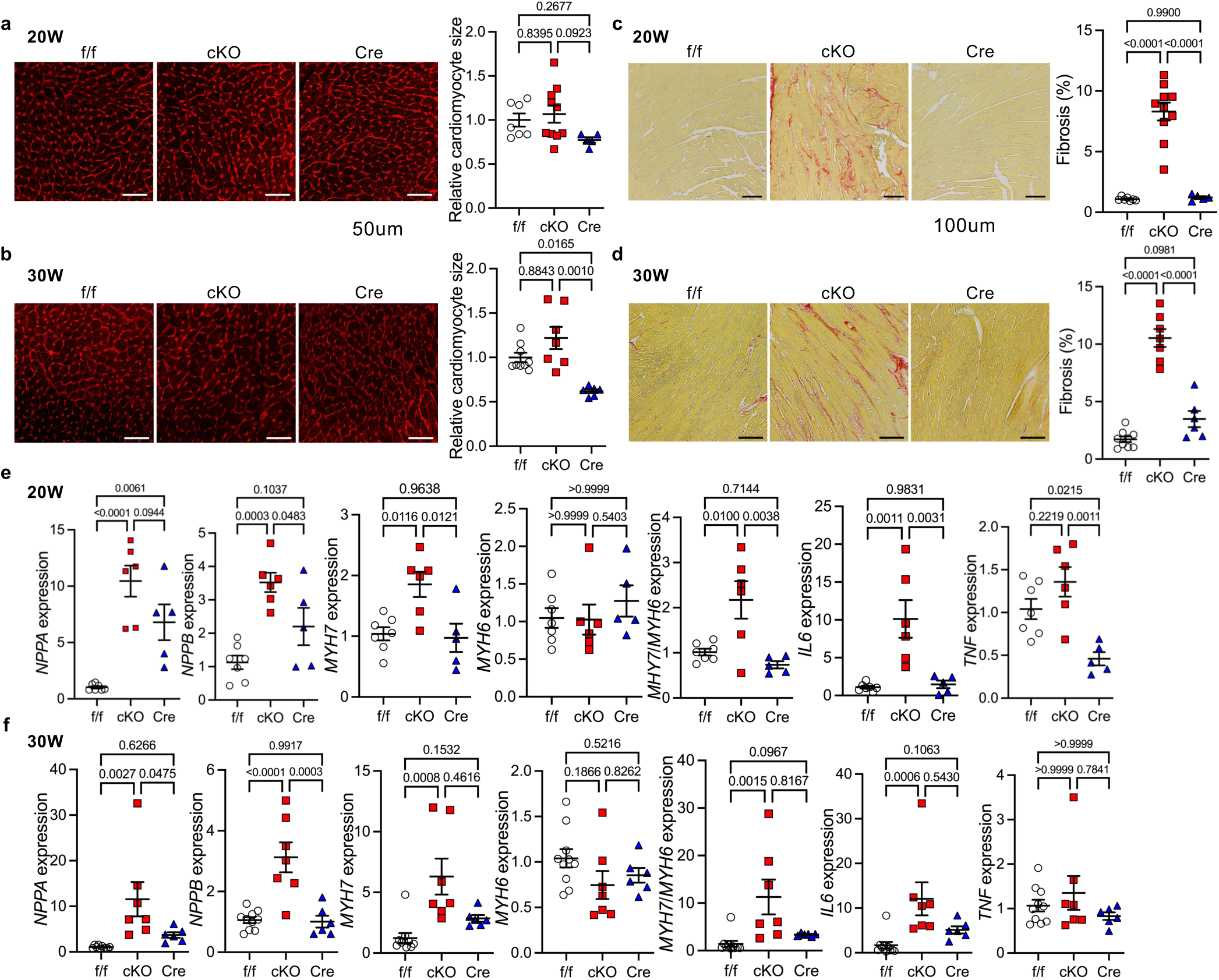
SCOT cardiac-specific deletion induces heart failure with severe cardiac inflammation in male mice. a-b, Wheat germ agglutinin (WGA) staining showing cardiomyocyte size, with quantitative analysis for 19-22-week-old (a) and 28-30-week-old (b) mice. **c-d**, Picric Acid Sirius Red (PASR) staining, indicating cardiac fibrosis, with quantitative analysis for 19-22-week-old (c) and 28-30-week-old (d) mice. **e-f**, Gene expression analysis of markers associated with heart failure, cardiac remodeling, and inflammation in 19-22-week-old (e) and 28-30-week-old (f) mouse hearts. n = 7 (f/f), 10 (cKO), and 5 (Cre) (a, c); n = 9 (f/f), 7 (cKO), and 6 (Cre) (b, d); n = 7 (f/f), 6 (cKO), and 5 (Cre) (e); n = 10 (f/f), 7 (cKO), and 6 (Cre) (f). One-way ANOVA followed by Tukey’s multiple comparison test. *N* represents biologically independent replicates. *p-*values are shown in the figure. Data are presented as mean ± SEM. Source data are provided as a Source Data file.

To further investigate the effects of SCOT deletion in the heart, we examined the mRNA expression of genes associated with wall stress, remodeling, and inflammation. The expression levels of atrial natriuretic peptide (*NPPA*) and B-type natriuretic peptide (*NPPB*), markers of ventricular wall stress and heart failure, were significantly elevated in SCOT cKO mice as early as 19-22 weeks of age (Fig. 3e, 3f). Similarly, *MYH7* expression and the *MYH7*/*MYH6* ratio, markers of cardiac remodeling, were also significantly increased at this time point in SCOT cKO mice (Fig. 3e, 3f). Additionally, consistent with the histological analysis, the gene expression of inflammatory cytokines, *IL6* and *TNF*, was upregulated in SCOT cKO mice as early as 19-22 weeks of age (Fig. 3e, 3f). These findings are consistent with the observation from echocardiographic and necropsy analyses, further supporting the roles of SCOT deletion in progressive cardiac dysfunction and remodeling in male mice.

### The effects of SCOT deletion on cardiac function and morphology are sex-independent

Cardiac metabolism differs between males and females due to sex hormones and genetic factors; however, the role of sex differences in SCOT-mediated ketone catabolism remains unexplored in the heart. To investigate this, we monitored female SCOT cKO mice for up to 30 weeks of age under a normal chow diet, performing echocardiographic, biochemical, necropsy, and histological analyses (Fig. 4, 4a, 4b). Similar to male SCOT cKO mice, no differences in cardiac function, as assessed by echocardiography, were observed in female SCOT cKO mice aged 10 weeks or younger (Fig. 4a, 4c). However, cardiac dilatation and systolic dysfunction became evident as early as 19-22 weeks of age in female SCOT cKO mice (Fig. 4a, 4d). Consistent with findings in male mice, LV posterior wall thickness in diastole was reduced as early as 19-22 weeks of age in female SCOT cKO mice (Fig. 4d, 4e). Necropsy analysis revealed no significant differences between groups except for ventricular weight, which increased after 28 weeks of age, coinciding with the onset of cardiac dysfunction and an associated increase in lung weight (Fig. 4f, 4fj). Gross appearance of the heart showed cardiac enlargement as early as 19 weeks of age (Fig 4i), indicating cardiac remodeling. Consistent with observations in male mice, cardiac SCOT deletion did not alter serum β-OHB levels throughout aging in females (Fig. 4g, 4k). Furthermore, despite cardiac dysfunction, SCOT cKO mice did not exhibit cardiomyocyte hypertrophy over time, as assessed by WGA staining (Fig. 4h, 4l). These findings suggest that SCOT-mediated ketone catabolism is essential for maintaining cardiac homeostasis during aging and that its disruption leads to contractile dysfunction and cardiac dilatation without overt hypertrophy. Moreover, the effects of SCOT downregulation on cardiac function and morphology appear to be more pronounced in females.

**Figure 4:**
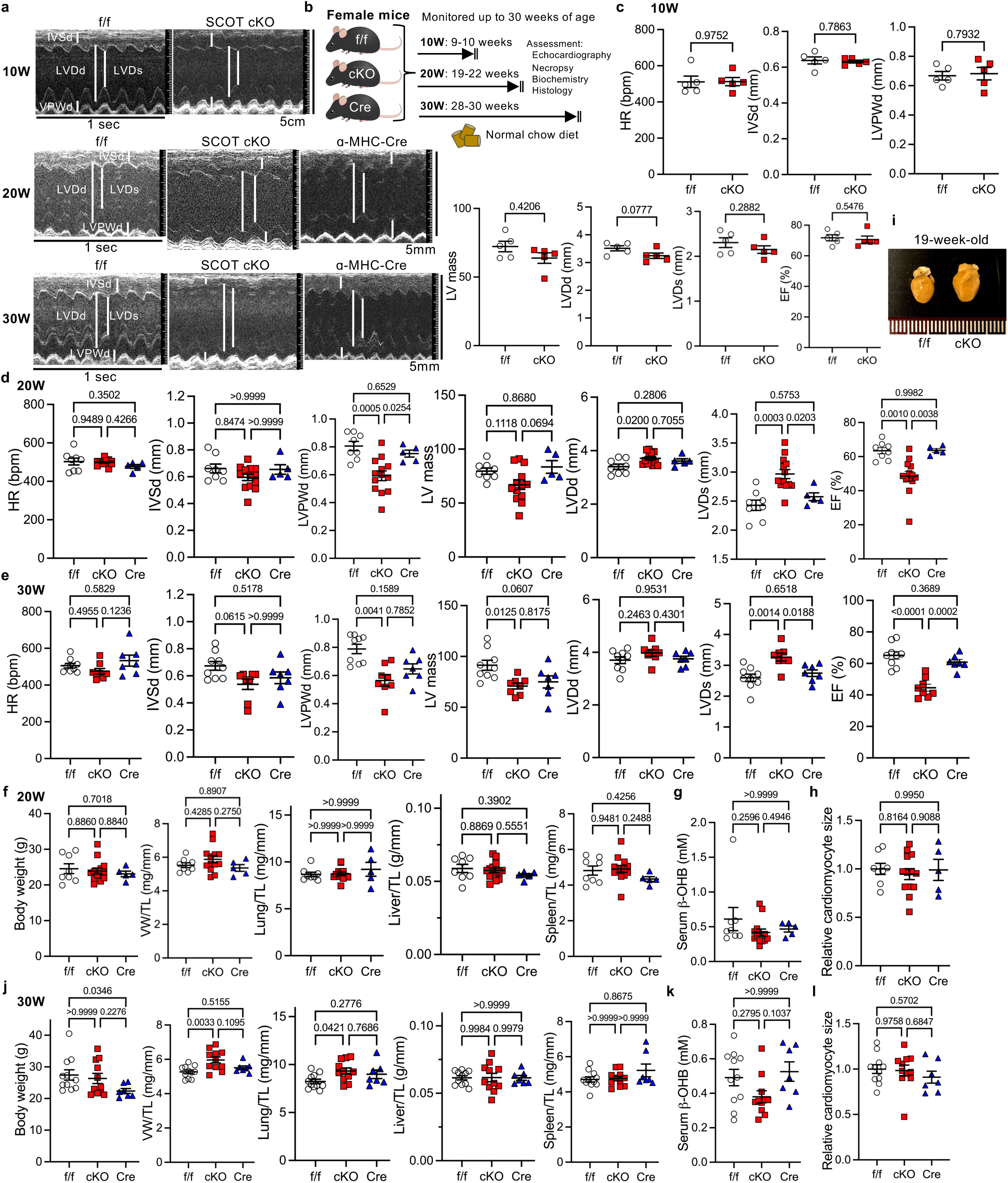
Cardiac-specific deletion of SCOT induces cardiac dysfunction in the absence of overt hypertrophy in female mice. **a-b**, Female mice were monitored up to 30 weeks of age. Representative M-mode echocardiography images (a). Vertical scale bar, 5mm; transverse scale bar, 1 sec. Schematic representation of the experimental design (b). **c-e**, Echocardiographic parameters of female mice at the indicated ages. n = 5 (c); n = 8 (f/f), 13 (cKO), and 5 (Cre) (d); n = 9 (f/f), 8 (cKO), and 7 (Cre) (e). **f-i**, Necropsy data of female mice at 19-22 weeks of age. n = 8 (f/f), 13 (cKO), and 5 (Cre). **j-l**, Necropsy data of female mice at 28-30 weeks of age. n = 11 (f/f), 11 (cKO), and 7 (Cre). Necropsy data including body weight, ventricular weight (VW) normalized to tibia length (TL), lung weight/TL, liver weight/TL, and spleen weight/TL (f, j). Serum β-hydroxybutyrate levels (g, k). Relative cardiomyocyte size, assessed by wheat germ agglutinin staining (h, l). Gross heart appearance of the indicated mice (i). Scale unit: mm. One-way ANOVA followed by Tukey’s multiple comparison test. *N* represents biologically independent replicates. *p* values are shown in the figure. Data are presented as mean ± SEM. Source data are provided as a Source Data file.

### A ketogenic diet affects systemic metabolism independently of cardiac SCOT expression in both sexes

Since SCOT is predominantly expressed in cardiomyocytes (Fig. 1d), we speculated that ketone supplementation may not directly enhance ketone oxidation in the hearts of SCOT cKO mice. Instead, the accumulation of ketone metabolites due to SCOT deletion could exert cardiotoxic effects, potentially through hyperacetylation of mitochondrial proteins or disruption of the TCA cycle by sequestering CoA in non-cardiomyocytes in the heart. To distinguish potential ketone metabolite toxicity from diminished ketone oxidation and to gain mechanistic insights into how SCOT deletion leads to systolic dysfunction and cardiac dilatation, we administered ketones through a ketogenic diet in SCOT cKO mice.

Ketones exert pleiotropic effects on metabolism and signaling pathways (25). To assess their systemic impact, we first examined how a short-term (12-day) ketogenic diet influences SCOT cKO and control f/f or αMHC-Cre alone mice (Fig. 5a). This early time point was specifically chosen to eliminate potential secondary effects of the ketogenic diet arising from cardiac dysfunction, allowing for a more precise evaluation of its systemic effects. Interestingly, the short-term ketogenic diet reduced body weight gain in male mice across both f/f and cKO groups, through the effect was not statistically significant, consistent with its well-documented metabolic effects, whereas it significantly increased body weight gain in female mice in both groups (Fig. 5b). Random glucose levels were decreased by the ketogenic diet in both sexes, with a more pronounced effect observed in females (Fig. 5c). Notably, even a short-term ketogenic diet significantly elevated serum β-OHB levels, reaching approximately twice the baseline levels in both sexes, regardless of SCOT expression levels in the heart (Fig. 5d). These findings indicate that ketone supplementation from a ketogenic diet has a systemic metabolic impact, while cardiac SCOT expression has no remarkable effect on systemic metabolism in response to short-term ketogenic diet exposure.

**Figure 5:**
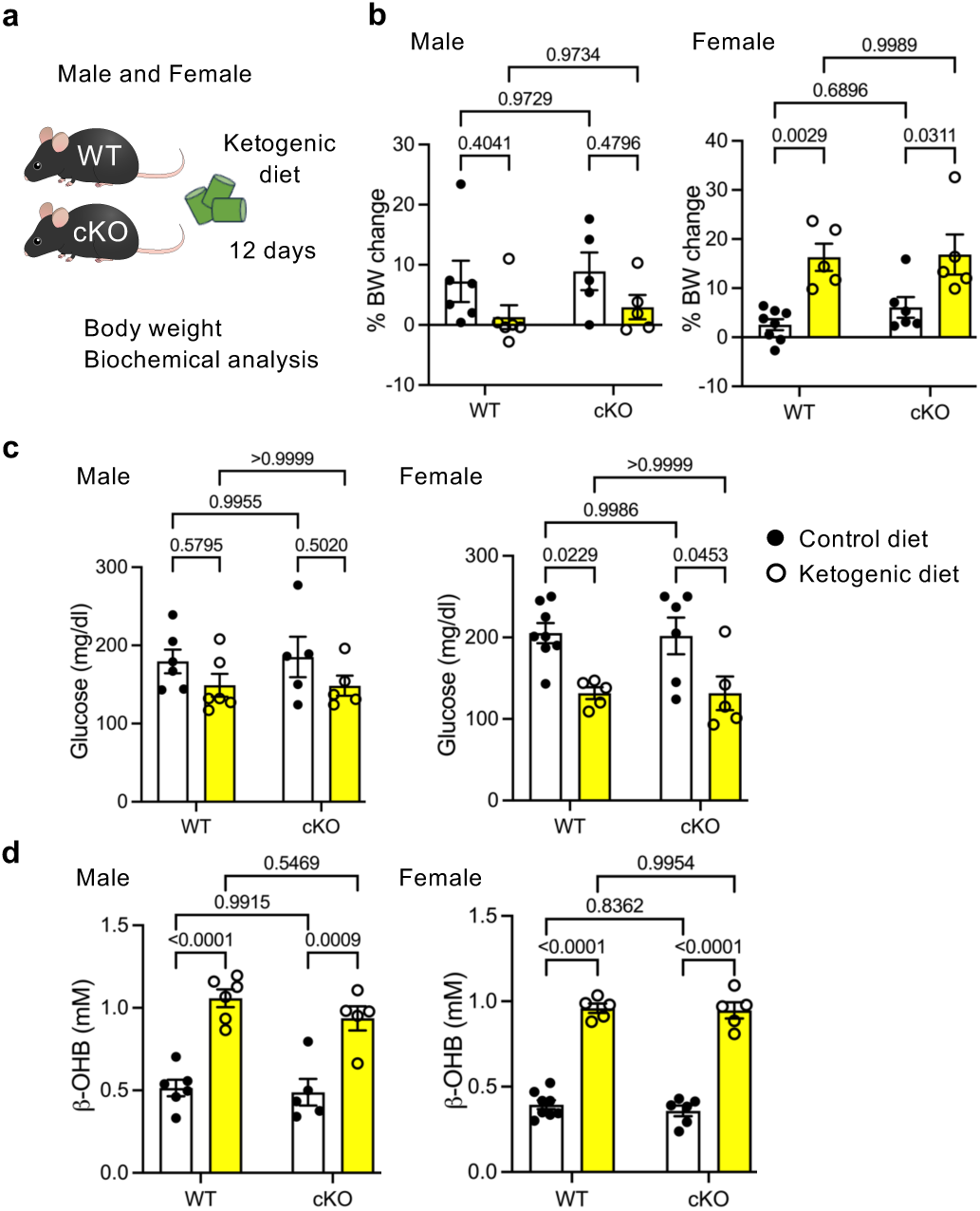
Systemic effects of a ketogenic diet in SCOT cardiac-specific knockout mice. a, Schematic representation of the experimental design. **b,** Body weight change in SCOT cardiac-specific knockout (cKO) and control (WT: flox/flox or αMHC-Cre alone) mice fed a ketogenic or control diet for 12 days, starting at 9-10 weeks of age. **c**, Casual plasma glucose levels. **d**, Serum β-hydroxybutyrate (β-OHB) levels. n = 6 (WT) and 5 (cKO) in male mice; n = 8 (WT) and 6 (cKO) in the control diet group and 5 in the ketogenic diet group for female mice. Two-way ANOVA followed by Tukey’s multiple comparison test. *N* represents biologically independent replicates. *p* values are indicated in the figure. Data are presented as mean ± SEM. Source data are provided as a Source Data file.

### A ketogenic diet partially attenuates SCOT deletion-induced cardiac dysfunction in male mice

Given that a ketogenic diet has been shown to attenuate pressure overload-induced heart failure in mice (22), we investigated how elevated serum ketone levels via a ketogenic diet impact SCOT-deletion-induced cardiac dysfunction. Control (f/f and αMHC-Cre alone) and SCOT cKO mice were placed on a ketogenic diet starting at 16 weeks of age for 4 weeks (Fig. 6a). The presence or absence of SCOT-mediated ketolysis did not affect body weight changes during ketogenic diet consumption (Fig. 6b). There were no differences in serum β-OHB levels among the groups after four weeks of ketogenic diet consumption (Fig. 6c). Cardiac function was then assessed by echocardiography (Fig. 6d). Although SCOT cKO mice already developed contractile dysfunction by 20 weeks of age, corresponding to the end of the ketogenic diet period, the ketogenic diet mitigated systolic dysfunction in SCOT cKO mice (Fig. 6e). Ventricular wall thickness and LV mass remained unchanged between groups. Necropsy findings supported the echocardiographic data, showing that heart, lung, and liver weights were not significantly affected by the ketogenic diet across all genotypes (Fig. 6f). However, spleen weight was slightly but significantly increased in αMHC-Cre mice (Fig. 6f). These findings suggest that a ketogenic diet confers cardioprotection even in the absence of SCOT-mediated ketolysis in the heart and that SCOT cKO mice develop contractile dysfunction and cardiac dilatation during aging, at least partially independent of reduced ketone oxidation.

**Figure 6:**
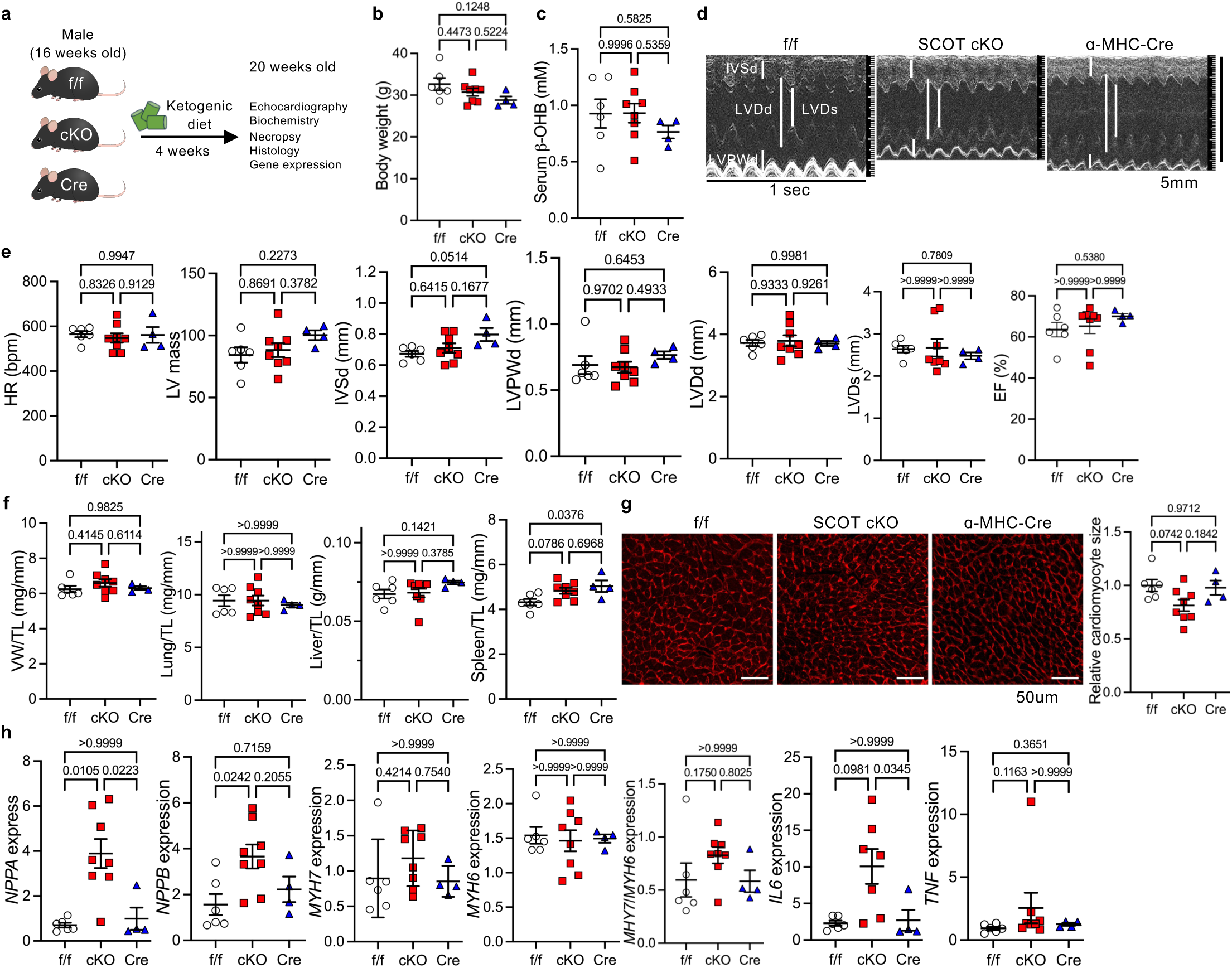
A ketogenic diet mitigates cardiomyopathy in cardiac-specific SCOT knockout male mice. a, Schematic representation of the experimental design. Data were collected after 4 weeks of ketogenic diet feeding. **b-c**, Body weight (b) and serum β-hydroxybutyrate (β-OHB) levels (c). **d-e**, Representative M-mode echocardiography images (d) and echocardiographic parameters (e). Vertical scale bar, 5mm; transverse scale bar, 1 sec. **f**, Necropsy data, including ventricular weight (VW) normalized to tibia length (TL), lung weight/TL, liver weight/TL and spleen weight/TL. **g**, Wheat germ agglutinin (WGA) staining showing cardiomyocyte size, with quantitative analysis. **h**, Gene expression analysis of markers associated with heart failure, cardiac remodeling, and inflammation. n = 6 (f/f), 8 (cKO), and 4 (Cre). One-way ANOVA followed by Tukey’s multiple comparison test. *N* represents biologically independent replicates. *p* values are shown in the figure. Data are presented as mean ± SEM. Source data are provided as a Source Data file.

Individual cardiomyocyte size remained comparable across all genotypes during ketogenic diet consumption (Fig. 6g), presumably due to the absence of cardiac hypertrophy at baseline in SCOT cKO mice. Gene expression analysis revealed that *NPPA* and *NPPB* expression remained elevated in SCOT cKO mouse hearts (Fig. 6h), suggesting that ventricular wall stress was not fully normalized by the ketogenic diet, despite the mitigation of contractile dysfunction. However, the expression of *MYH7* and inflammatory cytokines *IL6* and *TNF* was slightly but significantly reduced following ketogenic diet intervention (Fig. 6h). These findings suggest that exogenous ketones confer cardioprotective effects through a mechanism distinct from enhancing ketone oxidation, emphasizing the crucial role of ketones as signaling molecules rather than merely serving as oxidative substrates.

### A ketogenic diet mitigates SCOT deletion-induced cardiac dysfunction and remodeling in female mice

Since SCOT-mediated ketone catabolism is essential for maintaining cardiac homeostasis during aging, particularly in females, we investigated whether a ketogenic diet provides cardioprotection in SCOT cKO female mice as well. SCOT cKO and two control groups were placed on a ketogenic diet starting at 16 weeks of age for 4 weeks (Fig. 7a). The presence or absence of cardiac SCOT expression did not affect body weight during the 4-week ketogenic diet regime (Fig. 7b). The effect of the ketogenic diet on serum β-OHB levels was similar across all groups (Fig. 7c). Although SCOT cKO female mice exhibited not only systolic dysfunction but also cardiac remodeling, including a thinner ventricular wall, by 20 weeks of age (Fig. 4), the ketogenic diet attenuated the progression of contractile dysfunction and cardiac remodeling, mirroring the effects observed in male mice (Fig. 7d, 7e).

**Figure 7:**
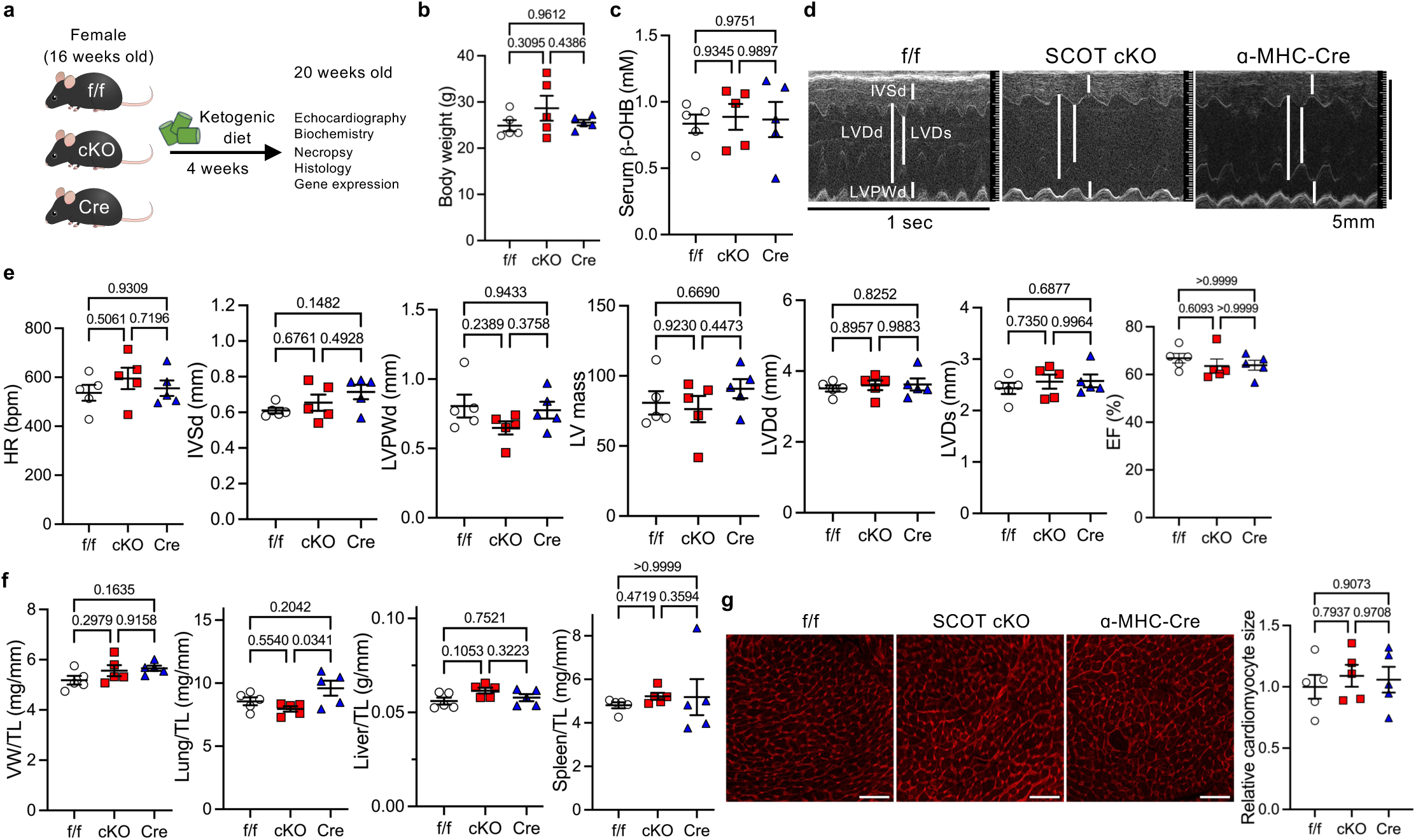
A ketogenic diet attenuates cardiomyopathy in cardiac-specific SCOT knockout female mice. a, Schematic representation of the experimental design. Data were collected after 4 weeks of ketogenic diet feeding. **b-c**, Body weight (b) and serum β-hydroxybutyrate (β-OHB) levels (c). **d-e**, Representative M-mode echocardiography images (d) and echocardiographic parameters (e). Vertical scale bar, 5mm; transverse scale bar, 1 sec. **f**, Necropsy data, including ventricular weight (VW) normalized to tibia length (TL), lung weight/TL, liver weight/TL and spleen weight/TL. **g**, Wheat germ agglutinin staining showing cardiomyocyte size, with quantitative analysis. n = 5. One-way ANOVA followed by Tukey’s multiple comparison test. *N* represents biologically independent replicates. *p* values are shown in the figure. Data are presented as mean ± SEM. Source data are provided as a Source Data file.

Consistent with echocardiographic data, ventricular weight normalized to tibia length was not significantly different between groups (Fig. 7f). Lung weight was slightly increased in αMHC-Cre mice compared to SCOT cKO mice (Fig. 7f). No differences in liver or spleen weights were observed among the three groups (Fig. 7f). Additionally, cardiomyocyte size remained unchanged regardless of SCOT expression during ketogenic diet consumption (Fig. 7g). These findings indicate that a ketogenic diet exerts cardioprotective effects primarily through modulation of signaling pathways rather than serving as an additional fuel source in the heart, irrespective of sex.

### SCOT upregulation alters mitochondrial function

Given that SCOT deletion induces cardiac contractile dysfunction and that ketone supplementation via a ketogenic diet mitigates cardiomyopathy caused by SCOT deletion in the heart, we hypothesized that SCOT expression influences mitochondrial function independently of ketone oxidation. To test this, we generated an adenovirus harboring FLAG-tagged SCOT (Ad-FLAG-SCOT) (Fig. 8a). H9C2 cardiomyoblasts were transfected with Ad-FLAG-SCOT for 2 days, followed by Seahorse experiments to assess mitochondrial respiration. As expected, SCOT overexpression did not affect basal oxygen consumption rate (OCR), ATP-linked respiration, or non-mitochondrial respiration under basal conditions (Fig. 8b, 8c). However, SCOT overexpression enhanced maximal respiratory capacity (Fig. 8b, 8c), suggesting that upregulation of SCOT positively impacts mitochondrial function, promoting long-term cellular survival and resistance to stress.

**Figure 8:**
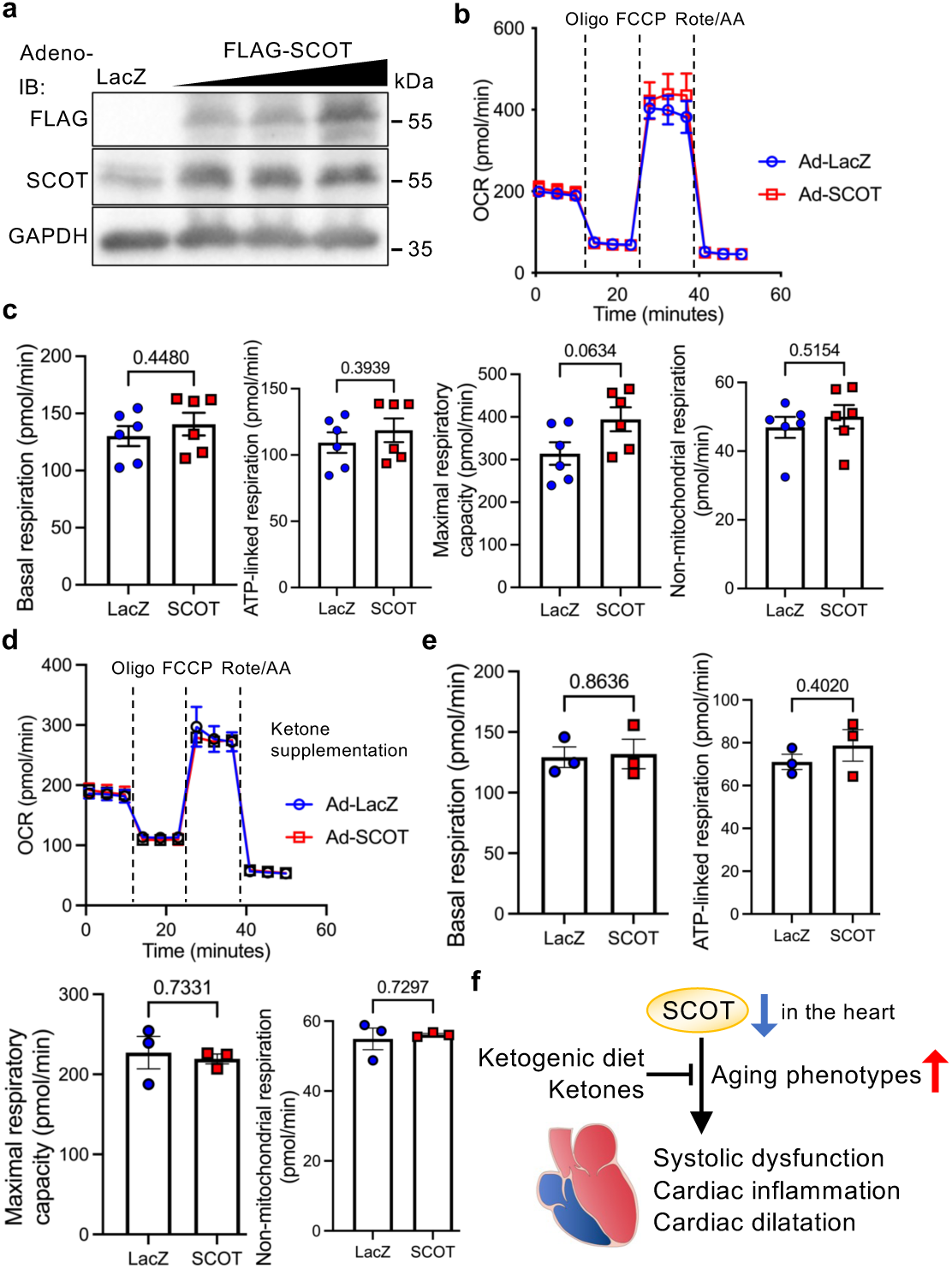
SCOT overexpression enhances mitochondrial maximal respiratory capacity, which is reversed by ketone supplementation in H9C2 cells. **a**, Immunoblots showing adenovirus-mediated FLAG-tagged SCOT overexpression in H9C2 cells. **b-e**, Mitochondrial function was assessed using Seahorse experiments. Representative oxygen consumption rate (OCR) plots in H9C2 cells transfected with FLAG-SCOT or control LacZ in the absence (b) or presence (d) of 100 µM β-hydroxybutyrate (β-OHB). Histograms showing basal respiration, ATP-linked respiration, maximal respiratory capacity, and non-mitochondrial respiration (c, e). n = 4 (b), 6 (c), 3 (d), and 3 (e). Unpaired t-test. *N* represents biologically independent replicates. *p* values are indicated in the figure. Data are presented as mean ± SEM. Source data are provided as a Source Data file.

Next, we conducted the same experiments in the presence of 100 µM β-OHB. Similar to conditions without ketones, SCOT overexpression did not affect mitochondrial respiratory function (Fig. 8d, 8e). However, unlike in the absence of ketones, ketone supplementation attenuated the enhanced maximal respiratory capacity induced by SCOT upregulation (Fig. 8d, 8e). These findings suggest that SCOT upregulation functions as a cardioprotective mechanism by enhancing mitochondrial maximal respiratory capacity independently of basal and ATP-linked respiration. Furthermore, ketones may normalize SCOT-mediated mitochondrial alterations, possibly through modulation of signaling pathways.

## DISCUSSION

In this study, we demonstrate that cardiac ketolysis is essential for maintaining structural and functional cardiac integrity during aging. Furthermore, ketone supplementation through a ketogenic diet mitigates SCOT deletion-induced cardiomyopathy. These findings suggest that a mechanism independent of ketone oxidation underlies both the development of cardiomyopathy in SCOT cKO mice and the cardioprotective effects of ketones (Fig. 8f).

Ketones serve as an alternative energy source for various organs, including the brain, heart, skeletal muscle, and kidneys, particularly during fasting and prolonged exercise (18). Decreased SCOT expression in the cerebellum and skeletal muscle has been reported in Friedreich’s ataxia, an inherited neurodegenerative disorder often accompanied by cardiomyopathy, where SCOT dysregulation contributes to neurological symptoms (27). Additionally, sensory neuron-specific SCOT KO mice, generated using an Advillin-Cre model, exhibited functional alterations in the somatosensory nervous system (28), suggesting that ketone metabolism plays a pivotal role in nervous system development and the progression of neurodegenerative diseases. However, in the heart, ketone oxidation contributes only a minor role in cardiac energy production, accounting for approximately 10-15% of ATP generation (2). Moreover, the heart exhibits metabolic flexibility, utilizing a variety of available nutrients. Based on this, we initially hypothesized that SCOT deletion would not significantly impact cardiac function. However, our findings reveal the opposite - ketone catabolism is essential for maintaining cardiac function throughout aging. This crucial role appears to stem not primarily from ketone oxidation contributing to cardiac energy production but rather from its function as a signaling mechanism.

Ketone utilization is highest during neonatal development and declines throughout adulthood in the nervous system, whereas in the heart, SCOT is highly expressed during adolescence and decreases with aging in female mice. In heart failure, ketone oxidation is increased in heart failure with reduced ejection fraction (HFrEF) but decreased in heart failure with preserved ejection fraction (HFpEF) and diabetic cardiomyopathy (29). These developmental stage-dependent, tissue-specific, and disease-specific changes in SCOT expression highlight the crucial role of ketolysis in maintaining organ homeostasis. Ketogenic diets or ketone esters have been shown to provide neuroprotection and cardioprotection against pediatric refractory epilepsy, seizures, and heart failure (22, 30–33), emphasizing the importance of ketones in preventing disease progression through distinct mechanisms depending on the disease type. It is importance to note that ketogenic diets and ketone esters likely exert cardioprotection via partially distinct pathways, as ketogenic diets contain a very high fat content, which enhances fatty acid oxidation and reduces glucose oxidation with the decreased expression of key ketolysis enzymes in the heart (22, 34). Our study revealed that a ketogenic diet alleviates cardiomyopathy in SCOT cKO mice, indicating that ketogenic diet-mediated cardioprotection occurs through a ketone oxidation-independent mechanism. Further investigation is warranted to elucidate the underlying mechanisms by which ketones confer cardioprotection.

Ischemic heart disease is a major cause of heart failure (35), although contractile dysfunction can also develop in the absence of ischemia, a condition collectively referred to as non-ischemic cardiomyopathy. The primary cause of non-ischemic cardiomyopathy is hypertensive heart disease; however, this is unlikely in SCOT cKO mice, given that SCOT is modulated in a cardiomyocyte-specific manner. Another common cause is metabolism-related cardiomyopathy, including diabetic and obesity cardiomyopathy (36–38). Since SCOT is an enzyme involved in ketone catabolism in the heart, SCOT downregulation-induced heart disease can be specifically classified as a form of metabolism-related cardiomyopathy. Although systemic metabolism remains normal, including body weight, ketone, and glucose levels, we propose that SCOT downregulation plays a crucial role in the development of aging-associated heart disease through metabolic disturbance, particularly in female mice, as SCOT expression declines with age. However, the molecular mechanism by which SCOT downregulation leads to contractile dysfunction remains unclear. While ketones can mitigate SCOT-related heart disease, further investigation is needed to elucidate the underlying molecular mechanisms as to how downregulation of ketone catabolism contributes to contractile dysfunction, presumably through ketolysis inhibition-induced alteration of signaling pathways.

Cardiac fibrosis is prominently observed in SCOT cKO mice (Fig. 3). While myocarditis can induce acute cardiac decompensation with systolic dysfunction, SCOT cardiac-specific deletion leads to a gradual decline in function. Additionally, inflammation is a key driver of hypertrophy and diastolic dysfunction (3), a common cause of diabetic and obesity cardiomyopathy (37). Thus, although fibrosis may contribute to exacerbating contractile function, it is unlikely to be the primary cause of systolic dysfunction in SCOT cKO mice. Immunoblot analysis (Fig. 1b) confirms that SCOT protein expression is nearly absent in cKO mouse hearts, indicating that SCOT is predominantly and almost exclusively expressed in cardiomyocytes. To sustain uninterrupted ATP production, cardiomyocytes contain a high mitochondrial content to support oxidative phosphorylation. Since SCOT overexpression enhances maximal respiratory capacity, its downregulation may plausibly contribute to a reduction in maximal respiration, potentially leading to mitochondrial dysfunction during aging or increasing susceptibility to age-related decline.

In conclusion, our study demonstrates that downregulation of ketolysis in the heart progressively induces contractile dysfunction and fibrosis during aging. Moreover, ketone supplementation through a ketogenic diet mitigates cardiomyopathy stemming from impaired ketolysis.

## METHODS

### Mice

The Oxct1 mouse strain was created from ES cell clone EPD0082_1_C02_M25, obtained from the KOMP Repository (www.komp.org) and generated by the Wellcome Trust Sanger Institute (WTSI). Targeting vectors used were generated by the Wellcome Trust Sanger Institute and the Children’s Hospital Oakland Research Institute as part of the Knockout Mouse Project (3U01HG004080) (39). Oxct1 floxed mice were generated and purchased from KOMP (KO-6867 Oxct1tm1c(KOMP)Wtsi). Genotyping was performed using a forward (5’-CCAGCATTTATAGTAGCATGGAAATC-3’) and reverse primers (5’-TCAAACCTCCACAAGTGGTACAGGG-3’) to identify the floxed mice. C57BL/6J wild-type mice were purchased from the Jackson Laboratory (Strain #: 000664) at 5-8 weeks of age. Littermate SCOT floxed mice and αMHC-Cre mice were used as a control. Mice of both sexes were used in this study. Mice were housed in a temperature and humidity-controlled environment within a range of 21°C - 23°C with 12-hour light/dark cycles, in which they received food and water *ad libitum*. The sample size required was estimated to be n = 5-9 per group according to the Power analysis based upon previous studies examining the effects of a ketogenic diet on cardiac hypertrophy and function. All protocols concerning the use of animals were approved by the Institutional Animal Care and Use Committee at Rutgers New Jersey Medical School and all procedures conformed to NIH guidelines (Guide for the Care and Use of Laboratory Animals). Handling of mice and euthanasia with CO_2_ in an appropriate chamber were conducted in accordance with guidelines on euthanasia of the American Veterinary Medical Association. Rutgers is accredited by AAALAC International, in compliance with Animal Welfare Act regulations and Public Health Service (PHS) Policy on Humane Care and Use of Laboratory Animals, and has a PHS Approved Animal Welfare Assurance with the NIH Office of Laboratory Animal Welfare (D16-00098 (A3158-01)).

### Sex as a biological variable

Our study examined the effects of SCOT deletion and ketogenic diets in both male and female mice, and sex-dimorphic effects are reported.

### Ketogenic diet

Mice were fed a custom diet (Ketogenic diet (D22021702, 7 kcal% Protein, and 90% kcal% Fat) or Control diet (D22021701, 10 kcal% Protein, and 15 kcal% Fat) (Supplementary Table 1), purchased from Research Diets) ad libitum. Daily *ad libitum* food intake was measured after an initial 2-day acclimation period on a special diet followed by 12-day or 4-week experimental periods by weighing food provided and remaining every 3 days and taking an average. The body weight was measured in the late afternoon every 3 days during the special food feeding period. Random blood sugar levels were measured using Aimstrip plus blood glucose meter kit (VWR, #10025-286).

### β-hydroxybutyrate measurements

Blood was collected from the ventricular cavity of the heart at the time of euthanasia, kept on ice for 30 minutes, and then centrifuged at 2,000 x g for 10 minutes. The resulting supernatant was stored at -80°C until analysis. Serum β-hydroxybutyrate levels were measured using a β-hydroxybutyrate colorimetric assay kit (Caymen #700190) following the manufacturer’s instructions.

### Echocardiography

Mice were anesthetized using 10 μl/g body weight of 2.5% avertin (Sigma-Aldrich), and echocardiography was performed using ultrasound (Vivid 7, GE Healthcare). It took around 10-20 minutes from the establishment of anesthesia to the completion of echocardiography and 1-2 hours to fully recover from anesthesia after echocardiography. A 13-MHz linear ultrasound transducer was used. Mice were subjected to 2-dimension guided M-mode measurements of LV internal diameter at the papillary muscle level from the short-axis view to measure heart rate, systolic function (ejection fraction with LV diameter at end diastole (LVDd) and LV diameter at end systole (LVDs)) and wall thickness (Interventricular septal at end diastole (IVSd)), which were taken from at least three beats and averaged. LV ejection fraction was calculated as follows: Ejection fraction = [(LVDd)^3^ – (LVESD)^3^]/(LVEDD)^3^ x 100. LV mass was calculated as follows: LV mass = 1.05 x [(IVSd+LVDd+LVPWd)^3^ – (LVDd)^3^].

### Cell line

HEK293 cells were purchased from the American Type Culture Collection (CRL-1573) and maintained at 37°C with 5% CO_2_ in Dulbecco’s modified Eagle’s Medium with 10% fetal bovine serum and penicillin/streptomycin. H9C2 cells were purchased from the ATCC and were maintained at 37 °C with 5% CO_2_ in Dulbecco’s modified Eagle’s medium/Nutrient Mixture F-12 supplemented with 10% fetal bovine serum and penicillin/streptomycin.

### Adenovirus construct

Recombinant adenovirus vector for overexpression was constructed, propagated and tittered as previously described (40–42). Briefly, pBHGloxΔE1,3Cre (Microbix), including the ΔE adenoviral genome, was co-transfected with the pDC shuttle vector containing the gene of interest into HEK293 cells. Mouse *OXCT1* cDNA was generated using mRNA isolated from mouse hearts. The *OXCT1* cDNA sequence was confirmed by Sanger Sequencing. Replication-defective human adenovirus type 5 (devoid of E1) harboring mouse full length *OXCT1* cDNA (Ad-FLAG-tagged OXCT1) was generated by homologous recombination in HEK293 cells. Adenovirus harboring beta-galactosidase (Ad-LacZ) was used as a control.

### Immunoblotting

Cardiomyocyte lysates and heart homogenates were prepared in RIPA with protease and phosphatase inhibitors (Sigma-Aldrich, P8340, P0044) as described previously (40). In shirt, lysates were centrifuged at 13,200 rpm at 4°C for 15 minutes. Protein concentrations were determined using a standard BCA assay (Thermo Fisher Scientific, 23227). Total protein lysates (25-45 μg) were incubated with SDS sample buffer (final concentration: 100 mM Tris (pH 6.8), 2% SDS, 5% glycerol, 2.5% 2mercaptoethanol, and 0.05% bromophenol blue) at 95°C for 5 minutes. The denatured protein samples were separated by SDS-PAGE, transferred to polyvinylidene difluoride membranes (Bio-Rad, 1620177) by semi-dry electrophoretic transfer (Bio-Rad), blocked in 2% (w/v) BSA (Fisher, BP9703100) in 1xTBS/0.5% Tween 20 at room temperature for 1 hour, and probed with primary antibodies at 4°C overnight. After washing with 1xTBS/0.5% Tween 20 for 15 minutes, the membranes were incubated with the corresponding secondary antibody at room temperature for 1 hour. After washing with 1xTBS/0.5% Tween 20 for 30 minutes, the membranes were developed with ECL Western blotting substrate (Millipore, WBKLS0500, WBLUC0100), followed by acquisition of digital image with the ChemiDoc MP Imaging System (Bio-Rad).

### Antibodies and reagents

The following commercial antibodies were used at the indicated dilutions: rabbit monoclonal GAPDH (#5174) (1:5,000), anti-mouse or -rabbit IgG, HRP-linked antibodies (#7076 and #7074) (1:5,000) (Cell Signaling Technology); mouse monoclonal α-tubulin antibody (T6074) (1:5,000), mouse monoclonal FLAG M2 antibody (F1804) (1:4,000) (Sigma-Aldrich); and mouse polyclonal anti-BDH1 (#ab68321) (1:3,500) and rabbit polyclonal anti-OXCT1 (#ab105320) (1:3,500) (Abcam). Antibodies were diluted in either 2% (w/v) BSA in 1xTBS/0.5% Tween 20 or Immunobooster solution (Takara, T7111A), depending on the level of background intensity.

### Quantitative RT-PCR

Total RNA was harvested from mouse tissues using TRIzol reagent (Thermo Fisher Scientific, 15596-018) according to the manufacturer’s instructions. Complementary DNA (cDNA) was synthesized by reverse transcription using 480 ng total RNA with PrimeScript RT Master Mix (Takara, RR036). Using Maxima SYBR Green qPCR master mix (ThermoFisher Scientific, K0253), real-time RT-PCR was performed Bio-Rad CFX 96 Real-Time Detection System (Bio-Rad) under the following conditions: denaturing at 95°C for 3 min, 40 cycles of denaturing at 95°C for 10 s, annealing at 58°C for 15 s, extension at 72°C for 45 s, and a final elongation step at 72°C for 10 min. Cq values were generated by CFX Manager Software 3.1 (#1845000; Bio-Rad). Relative mRNA expression was determined by the ΔΔ-Ct method normalized to the ribosomal RNA (18S) level. The following oligonucleotide primers were used: NPPA, sense 5’-ATGGGCTCCTTCTCCATCAC-3’ and antisense 5’-ATCTTCGGTACCGGAAGCTG-3’; NPPB, sense 5’-AAGTCCTAGCCAGTCTCCAGA-3’ and antisense 5’-GAGCTGTCTCTGGGCCATTTC-3’; MYH7, sense 5’-GCCAACACCAACCTGTCCAAGTTC-3’ and antisense 5’-TGCAAAGGCTCCAGGTCTGAGGGC-3’; MYH6, sense 5’-GGAAGAGTGAGCGGCGCATCAAGG-3’ and antisense 5’-CTGCTGGAGAGGTTATTCCTCG-3’; IL6, sense 5’-CCGGAGAGGAGACTTCACAG-3’ and antisense 5’-TGCCATTGCACAACTCTTTT-3’; TNF, sense 5’-CCCTCACACTCAGATCATCTTCT-3’ and antisense 5’-GCTACGACGTGGGCTACAG-3’; 18S rRNA, sense 5’-CGCGGTTCTATTTTGTTGGT-3’ and antisense 5’-AGTCGGCATCGTTTATGGTC-3’.

### Immunohistochemistory

Heart specimens were fixed with formalin, embedded in paraffin, and sectioned at (6) µm thickness. Interstitial fibrosis was evaluated by Masson’s Trichrome and Picric Acid Sirius Red (PASR) staining. The myocyte cross-sectional area was measured from images captured from wheat germ agglutinin (WGA)-stained sections as described previously (43). The tissues were observed under a fluorescence microscope (BX51, Olympus). Images were extracted using the cellSens software (cellSens Standard, software version 4.2, Olympus). The outlines of 100–200 myocytes were traced in each section using NIH ImageJ.

### Seahorse experiments

The basal oxygen consumption rate (OCR) in H9C2 cells was measured by a Seahorse XFe96 Extracellular Flux Analyzer (Seahorse Bioscience). H9C2 cells were plated in 6-well plates, followed by transduction with Adenovirus for 48 hours at 37°C with 5% CO_2_ in Dulbecco’s modified Eagle’s medium (DMEM) supplemented with 10% fetal bovine serum and penicillin/streptomycin prior to measurement. The next day, cells were plated at a density of 30,000 cells/well in 96-well Seahorse assay plates except for background correction wells (Agilent, 103794-100). One hour prior to the beginning of measurements, the medium was replaced with XF base medium (Agilent, 103335-100) supplemented with 5.5 mM glucose (Sigma-Aldrich, G8270), 1 mM pyruvate (Agilent, 103578-100) and 2 mM L-glutamine (Agilent, 103579-1001) and incubated for 1 hour in a 37°C incubator without CO_2_. Oxygen consumption rate (OCR) was measured three times at baseline, followed by injection with 10 µM oligomycin (Sigma-Aldrich, O4876) to measure the ATP-linked OCR. 5 µM Carbonylcyanidep-triflouromethoxyphenylhydrazone (FCCP) (Sigma-Aldrich, C2920), an uncoupler, was used to determine maximal respiration, and 5 μM rotenone (Sigma-Aldrich, R8875) and 5 μM antimycin A (Sigma-Aldrich, A8674) were injected to determine the non-mitochondrial respiration.

## STATISTICAL ANALYSIS

Data are represented as single data points or dot plots and all values are expressed as mean ± SEM. Statistical analyses were carried out by 2-tailed unpaired Student *t* test for 2 groups or one-way analysis of variance (ANOVA) followed by the Tukey post-hoc analysis for 3 groups or more unless otherwise stated. If the data distribution failed normality by the Shapiro–Wilk test or Kolmogorov–Smirnov test, the Mann–Whitney *U* test for 2 groups or Kruskal–Wallis test with the Dunn’s multiple comparison test for 3 groups or more was performed. Multiple group comparisons with time-course were performed by two-way ANOVA and the Tukey post-hoc analysis. The statistical analyses used for each figure are indicated in the corresponding figure legends. No statistical methods were used to predetermine sample size. The sample size was estimated based on data from published studies and pilot experiments. Investigators were blinded to data collection, measurement, and analysis. Owing to the nature of the cell culture experiments, randomization of the cell culture samples was not applicable. All experiments are represented by multiple biological replicates or independent experiments. The number of replicates per experiment are indicated in the legends. All experiments were conducted using at least two independent experimental materials or cohorts to reproduce similar results. No sample was excluded from analysis. GraphPad Prism 10 was used for statistical analysis and data visualization. A P-value of < 0.05 was considered significant.

## DATA AVAILABILITY

All data generated or analyzed during this study are included in this published article and its Supplementary Information. No original code was generated in this study. Any additional information required to reanalyze the data reported in this paper, reagents, and other materials are available from the corresponding author upon request.

## ACKNOWLEDGEMENTS

We thank Dr. Junichi Sadoshima, Chairman of the Department of Cell Biology and Molecular Medicine at Rutgers University, for his support. This study is supported in part by U.S. Public Health Service grants HL155766 (M.N.). This work was also supported by American Heart Association Scientist Development Grant (17SDG33660358) (M.N.).

## AUTHOR CONTRIBUTION

M.N. designed the experiments and wrote the paper; M.A.K and M.N. conducted the *in vitro* and *in vivo* experiments; M.A.K and M.N. conducted the animal experiments and analyses; M.A.K and M.N. wrote the paper. M.N. generated project resources. All authors reviewed and commented on the manuscript.

## FIGURE LEGENDS

**Supplementary Table 1:** The components of custom ketogenic diet and its isocaloric control diet, purchased from Research Diets, Inc.

| Diets (Product #) | D22021701 |  | D22021702 |  |
| --- | --- | --- | --- | --- |
|  | Control |  | Ketogenic |  |
|  | gm% | kcal% | gm% | kcal% |
| Protein | 10 | 10 | 12 | 7 |
| Carbohydrate | 74 | 75 | 5 | 3 |
| Fat | 7 | 15 | 67 | 90 |
| Total |  | 100 |  | 100 |
| Kcal/gm | 4.0 |  | 6.7 |  |
| Ingredient | gm | kcal | gm | kcal |
| Casein, 80 Mesh | 100 | 400 | 70 | 280 |
| L-Cystine | 1.5 | 6 | 1 | 4 |
| Corn Starch | 447.5 | 1790 | 0 | 0 |
| Maltodextrin 10 | 150 | 600 | 21.5 | 86 |
| Sucrose | 150 | 600 | 0 | 0 |
| Cellulose, BM200 | 50 | 0 | 50 | 0 |
| Soybean Oil | 25 | 225 | 25 | 225 |
| Lard | 0 | 0 | 0 | 0 |
| Corn Oil | 0 | 0 | 50 | 450 |
| Cocoa Butter | 0 | 0 | 50 | 450 |
| Hydrogenated Vegetable Oil | 44 | 396 | 280.2 | 2522 |
| Mineral Mix, S10026 | 10 | 0 | 10 | 0 |
| DiCalcium Phosphate | 13 | 0 | 13 | 0 |
| Calcium Carbonate | 5.5 | 0 | 5.5 | 0 |
| Potassium Citrate, 1 H<sub>2</sub>O | 16.5 | 0 | 16.5 | 0 |
| Vitamin Mix, V10001 | 10 | 40 | 10 | 40 |
| Choline Bitartrate | 2 | 0 | 2 | 0 |
| FD&C Yellow Dye #5 | 0.025 | 0 | 0.05 | 0 |
| FD&C Red Dye #40 | 0 | 0 | 0 | 0 |
| FD&C Blue Dye #1 | 0.025 | 0 | 0 | 0 |
| Total | 1025.05 | 4057 | 604.75 | 4057 |

